# New insights into the ORF2 capsid protein, a key player of the hepatitis E virus lifecycle

**DOI:** 10.1101/435933

**Authors:** Maliki Ankavay, Claire Montpellier, Ibrahim M. Sayed, Jean-Michel Saliou, Czeslaw Wychowski, Laure Saas, Sandrine Duvet, Cécile-Marie Aliouat-Denis, Rayan Farhat, Valentin de Masson d’Autume, Philip Meuleman, Jean Dubuisson, Laurence Cocquerel

## Abstract

Hepatitis E Virus (HEV) genome encodes three proteins including the ORF2 protein that is the viral capsid protein. Recently, we developed an efficient HEV cell culture system and demonstrated that this virus produces three different forms of its capsid protein: (i) the ORF2i form (infectious/intracellular) which is the form associated with the infectious particles, (ii) the ORF2g (glycosylated ORF2) and ORF2c (cleaved ORF2) forms that are massively secreted glycoproteins not associated with infectious particles, but are the major antigens present in HEV-infected patient sera. The ORF2 protein sequence contains three highly conserved potential N-glycosylation sites (N1, N2 and N3). Although ORF2 protein is the most characterized viral protein, its glycosylation status and the biological relevance of this post-translational modification is still unclear. In the present study, we constructed and extensively characterized a series of ORF2 mutants in which the three N-glycosylation sites were mutated individually or in combination. We demonstrated that the ORF2g/c protein is N-glycosylated on N1 and N3 sites but not on the N2 site. We showed that N-glycosylation of ORF2 protein does not play any role in replication and assembly of infectious HEV particles. We found that glycosylated ORF2g/c forms are very stable proteins which are targeted by patient antibodies. During our study, we also demonstrated that the ORF2i protein is translocated into the nucleus of infected cells. In conclusion, our study led to new insights into the molecular mechanisms of ORF2 expression.

## Importance

Hepatitis E virus (HEV) infection is the most common cause of acute viral hepatitis worldwide. This infection can become chronic in immunosuppressed patients and cause death in some patients. In our study, we focused on ORF2 viral capsid protein for which we recently identified glycosylated and non-glycosylated forms. The non-glycosylated form, named ORF2i, is the form associated with the infectious particles. The glycosylated forms, named ORF2g and ORF2c, are the major antigens present in HEV-infected patient sera. Here, we identified which sites on ORF2g/c proteins are N-glycosylated. We showed that N-glycosylation of ORF2 proteins is not involved in the biogenesis of infectious HEV particles. We found that ORF2g/c proteins are very stable and are targeted by patient antibodies. We also demonstrated that the ORF2i protein is translocated into the nucleus of infected cells. Thus, our study led to new important insights into the molecular mechanisms of ORF2 expression.

## Introduction

Hepatitis E virus (HEV) is the most common cause of acute viral hepatitis worldwide. This virus infects approximately 20 million people every year and is responsible for 3.4 million symptomatic cases and 70,000 deaths, mainly in regions of the world with low sanitary conditions (1). Although HEV infection is usually asymptomatic and self-resolving, severe forms in pregnant women (2) and chronic infections in immunocompromised patients (3) have been described. In addition, HEV infection has been associated with a broad range of extrahepatic manifestations, including renal and neurological manifestations (3). HEV strains infecting humans have been classified into 4 main distinct genotypes (gt) belonging to a single serotype. Gt1 and gt2 exclusively infect humans, are spread mainly through contaminated drinking water and represent main causes of waterborne outbreaks of hepatitis in developing countries. In contrast, gt3 and gt4 are zoonotic and mainly infect mammals. Their main reservoirs are pigs and game (4). The major transmission routes of gt3 and gt4 are direct contact with infected animals, consumption of contaminated food and transfusion of blood products. In industrialized countries, the most common genotype causing HEV infection is gt3. Importantly, due to the evolution toward chronicity in immunocompromised infected patients, HEV transmission through blood transfusion, resistance of some infected patients to ribavirin and complications in patients with preexisting liver disease, HEV infection is now considered as an emerging problem in industrialized countries (5).

HEV is a quasi-enveloped virus (6–8) containing a linear, single-stranded, positive-sense RNA genome that contains three open reading frames (ORFs), namely, ORF1, ORF2 and ORF3 (9). ORF1 encodes a non-structural polyprotein (ORF1 protein) that is essential for viral replication (10). It includes methyltransferase (Met), papain-like cysteine protease (PCP), RNA helicase (Hel) and RNA-dependent RNA polymerase (RdRp) domains (reviewed in (1)). ORF2 encodes the ORF2 viral capsid protein and ORF3 encodes a small multifunctional phosphoprotein that is involved in virion morphogenesis and egress (reviewed in (11)).

The ORF2 protein sequence contains 660 amino acids, a signal peptide sequence at its N-terminus and three highly conserved potential N-glycosylation sites represented by the sequon Asn-X-Ser/Thr (N-X-S/T) (12–14). The ORF2 protein has been largely studied and is the most characterized HEV viral protein. However, whether using various heterologous expression systems (14–17) or infectious systems (8, 13), the glycosylation status of ORF2 and the biological relevance of this post-translational modification was still controversial until recently. Indeed, by combining the highly replicative and cell culture-selected gt3 p6 strain (18) and a highly transfectable subclone of PLC/PRF/5 cells (PLC3 cells), we recently developed an efficient HEV cell culture system and demonstrated for the first time that, during its lifecycle, HEV produces 3 forms of the ORF2 capsid protein (**Figure 1A**): infectious/intracellular ORF2 (ORF2i), glycosylated ORF2 (ORF2g), and cleaved ORF2 (ORF2c) (19). We identified the precise sequence of the ORF2i and ORF2g proteins. The ORF2i protein is the form that is associated with infectious particles. The ORF2i protein is not glycosylated and is likely not translocated into the endoplasmic reticulum (ER) lumen and stays in the cytosolic compartment. In contrast, ORF2g and ORF2c proteins are secreted in large amounts in cell culture and infected patient sera, sialylated, N-and O-glycosylated but are not associated with infectious virions. The identification of these 3 forms of ORF2 led us to suggest the existence of two production pathways for the HEV capsid protein: (i) a major non-productive pathway in which ORF2 proteins are addressed to the secretion route where they are glycosylated, maturated and quickly secreted. (ii) a productive pathway in which cytosolic ORF2 proteins are delivered to the virion assembly sites (19). Importantly, ORF2g and ORF2c are the most abundant antigens detected in sera from patients (19). Whether ORF2g and ORF2c proteins play a specific role in the HEV lifecycle or have functions in host immune response needs to be elucidated.

**Figure 1:**
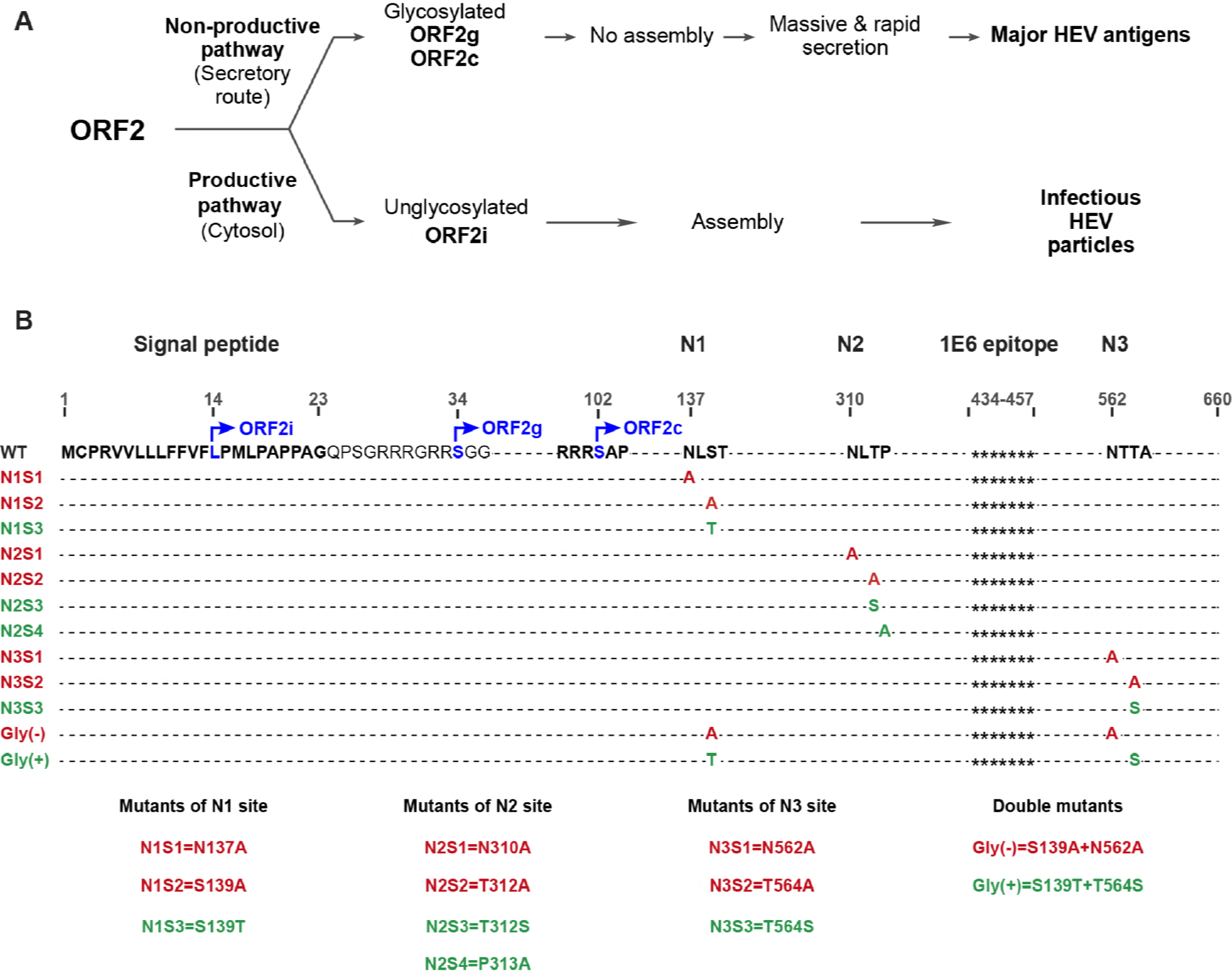
ORF2 production pathways and schematic representation of wild type and mutant ORF2 protein sequences. (**A**) ORF2 production pathways. The ORF2 protein follows two different pathways in the host cell (i) a major non-productive pathway in which ORF2 proteins are addressed to the secretion route where they are glycosylated, maturated and quickly secreted. This pathway leads to the production of the ORF2g and ORF2c proteins that are the major viral antigens present in the serum of HEV-infected patients. ORF2g and ORF2c proteins are N-glycosylated, O-glycosylated and sialylated. (ii) a productive pathway in which cytosolic ORF2i (infectious/intracellular) proteins are delivered to the virion assembly sites. The ORF2i protein is not glycosylated and is associated with infectious particles. (**B**) Schematic representation of wild type and mutant ORF2 protein sequences. HEV ORF2 protein is a 660 amino acid protein. The first 23 amino acids corresponding to the signal peptide are in bold. Positions of the first aa of ORF2i, ORF2g and ORF2c proteins are indicated. The three potential N-glycosylation sites are in bold (N1, N2 and N3). For each mutant, the introduced mutation(s) is/are shown. Mutations that inhibit N-glycosylation are in red whereas mutations that do not inhibit N-glycosylation are in green. The stars (*******) represent the epitope of the 1E6 anti-ORF2 antibody.

In the present study, we took advantage of our HEV cell culture system, in which ORF2 proteins are robustly expressed, to analyze the significance of N-glycosylation of the ORF2 protein in the HEV lifecycle. Using site-directed mutagenesis of the full-length infectious p6 clone, we constructed a series of ORF2 mutants in which the three potential N-glycosylation sites (^137^NLS, N1; ^310^NLT, N2; ^562^NTT, N3) were mutated individually or in combination. We performed an extensive characterization of these mutants by analyzing their subcellular localization, expression, oligomerization, stability and recognition by antibodies. We also studied the impact of mutations on assembly, density and infectivity of HEV particles. In addition to analyzing the glycosylation status of ORF2 protein and the biological relevance of this modification in the HEV life cycle, we obtained new insights into the molecular mechanisms of ORF2 expression.

## Results

### Generation of HEV p6 genomes expressing mutations of ORF2 N-glycosylation sites

The ORF2 protein sequence displays a signal peptide sequence at its N-terminus and three potential N-glycosylation sites, ^137^NLS (N1), ^310^NLT (N2), and ^562^NTT (N3) (**Figure 1B**). Previously, we demonstrated that the first amino acids (aa) of ORF2i and ORF2g proteins are Leu^14^ and Ser^34^, respectively (19). Here, by using the same mass spectrometry approach, we found that the first aa of ORF2c protein corresponds to Ser^102^ (**Figure 1B** and **Figure 2**), indicating that ORF2c protein is likely a cleavage product of the ORF2g protein.

**Figure 2:**
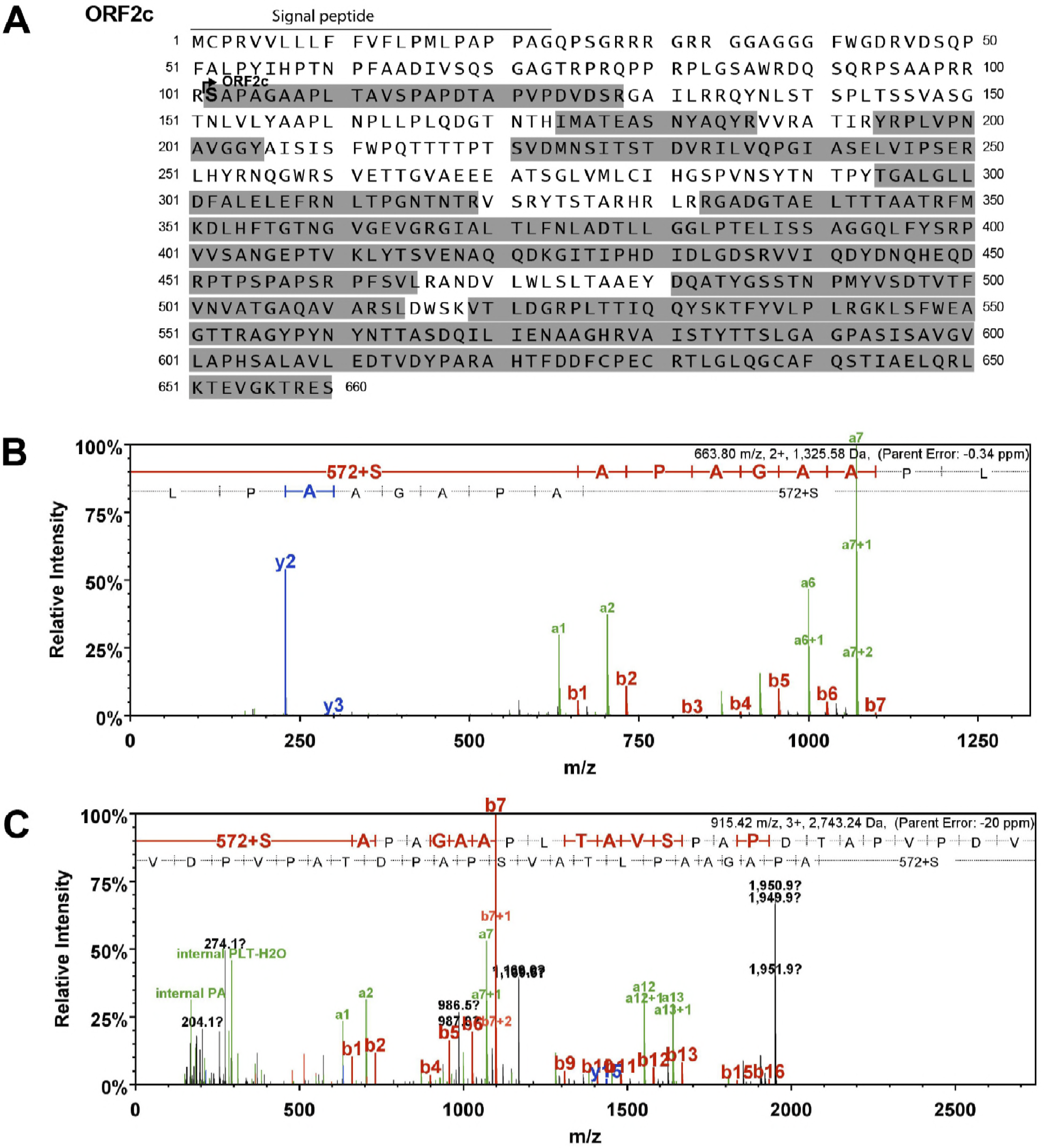
Identification of the N-terminus of the ORF2c protein. ORF2c proteins were immunoprecipitated with an anti-ORF2 antibody (4B2), denaturated and incubated or not with N-succinimidyloxycarbonylmethyl tris (2,4,6-trimethoxyphenyl) phosphonium bromide (TMPP), which binds specifically to the N-terminus of intact proteins. Proteins were resolved by SDS-PAGE, digested in-gel with trypsin or AspN and analyzed by nanoLC-MS/MS. (A) Peptide covering is highlighted in grey on the sequence. Ser^102^ in bold corresponds to the first aa of ORF2c that was identified by TMPP labeling. (B) and (C) MS/MS spectrum of N-terminal peptides of the ORF2c protein. (B) Tryptic peptide obtained from TMPP-labeled ORF2c protein. (C) AspN peptide obtained from TMPP-labeled ORF2c protein. +572 corresponds to the TMPP mass increment following TMPP labeling.

Using site-directed mutagenesis of the full-length infectious p6 clone, we constructed a series of ORF2 mutants in which N1, N2 and N3 sites were mutated individually (N1, N2 and N3 mutants) or in combination (Gly(−) and Gly(+) mutants) (**Figure 1B**). For each single mutant, three substitutions of the N-X-S/T sequon were generated: N-to-A (S1), S/T-to-A (S2) and S-to-T/T-to-S (S3). S1 and S2 substitutions were made to prevent N-glycosylation (Red), whereas S3 substitution does not affect N-glycan addition (Green). A fourth substitution (S4) was done for the N2 site in which a Proline (P) residue downstream of the sequon was replaced by an Alanine (^313^P-to-A). In Gly(−) and Gly(+) mutants, two combinations of mutations affecting (N1S2/N3S1) or not (N1S3/N3S3) the N-glycan addition on N1 and N3 glycosylation sites were introduced, respectively (**Figure 1B**). Mutants expressing mutations that theoritically abolish N-glycosylation are in Red. Mutants expressing mutations that theoritically do not disturb or improve N-glycosylation (N2S4) are in Green. Capped RNAs transcripts of wild type (wt) and mutant (mut) HEV genomes were generated and delivered into PLC3 cells by electroporation.

### Expression and subcellular localization of mutant ORF2 proteins

We first evaluated the expression and subcellular localization of mutant ORF2 proteins by indirect immunofluorescence. PLC3 cells electroporated with wild type and mutants of HEV-p6 RNAs (wt and mt PLC3/HEV-p6 cells) were fixed at 3 days post-electroporation (d.p.e.) and processed for ORF2 staining. For all constructs, over 90% of cells were ORF2-positive indicating that robust replication and expression of viral genome occurred in wt and mt PLC3/HEV-p6 cells (data not shown). As shown in **Figure 3A**, the wt ORF2 protein displayed mostly a cytoplasmic localization but also a nuclear localization, as recently described (20). Although mutant ORF2 proteins were globally expressed as the wt ORF2 protein, N1 mutants showed a slightly more perinuclear localization (**Figure 3A**), N2 mutants showed a concentrated staining in a spot close to the nucleus (**Figure 3B**) whereas N3 mutants showed a subcellular localization similar to that of wt ORF2 proteins (**Figure 3C**). Double labeling with anti-ORF2 and ER-specific anti-calnexin MAbs or with anti-ORF2 and Golgi-specific anti-GM130 MAbs failed to reveal co-localization of ORF2 proteins with these compartment markers (data not shown), as previously observed for the gt1 ORF2 protein (13). Nuclear localization of wt and mt ORF2 proteins was further characterized, as described below.

**Figure 3:**
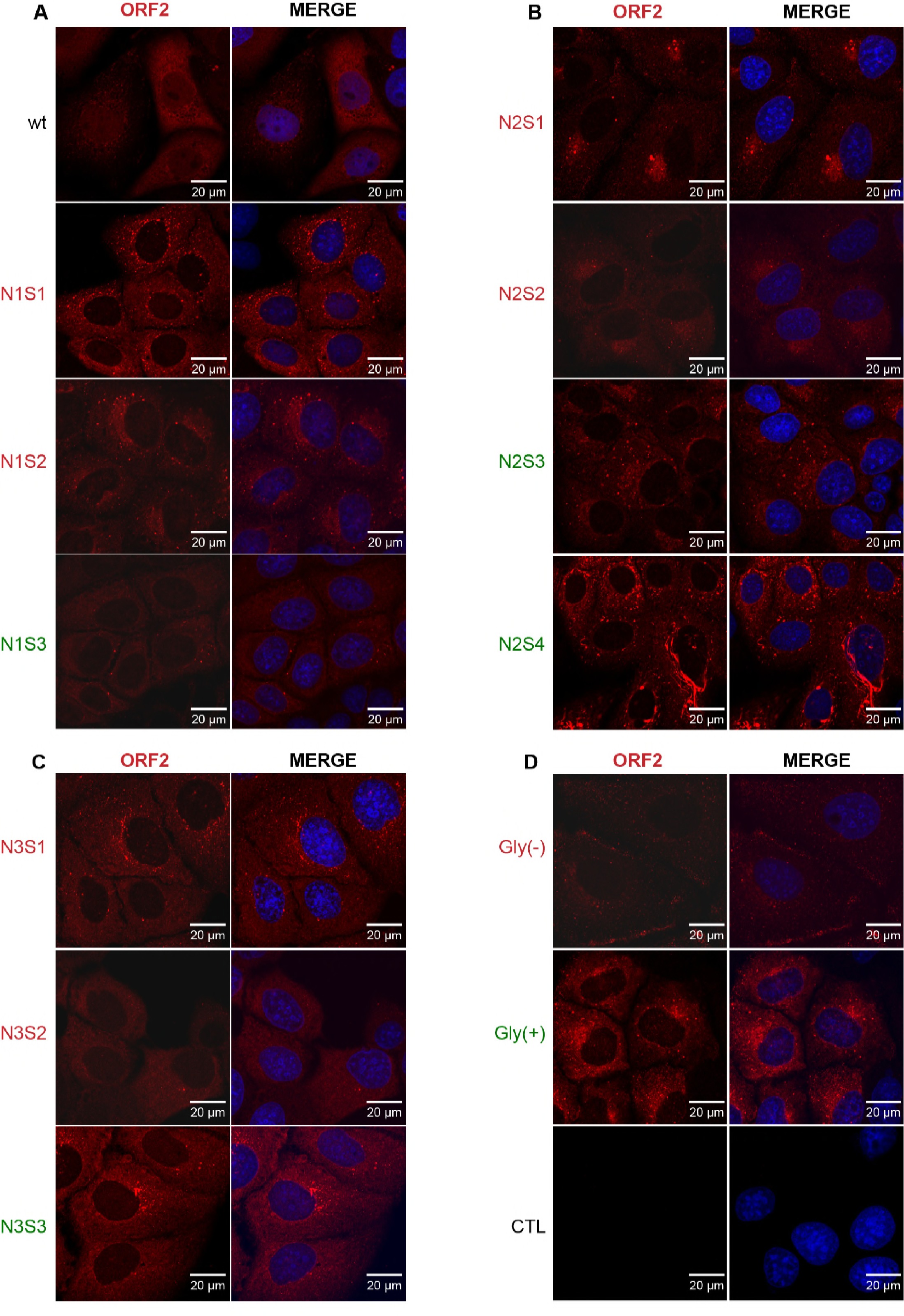
Subcellular distribution of wt and mutant ORF2 proteins. PLC3 cells were electroporated with wt and mt HEV-p6 RNAs and grown on coverslips. 3 days post-electroporation, cells were fixed and ORF2 protein was stained using the 1E6 antibody (in red). Nuclei were stained with DAPI (in blue). Cells were analyzed by confocal microscopy (magnification x40). Representative acquisitions of two independent experiments are presented. **(A)** Cells expressing wild type (wt) and N1S1 (N137A); N1S2 (S139A); N1S3 (S139T) mutants. **(B)** Cells expressing N2S1 (N310A); N2S2 (T312A); N2S3 (T312S) and N2S4 (P313A) mutants. **(C)** Cells expressing N3S1 (N562A); N3S2 (T564A); N3S3 (T564S)mutants. **(D)** Cells expressing Gly(−) (N1S2+N3S1) and Gly (+) (N1S3+N3S3) double mutants and non-transfected PLC3 cells (CTL).

Altogether, our results show that all the mutants of N-glycosylation sites of ORF2 protein are well expressed in PLC3 cells. Also, the wt ORF2 protein shows a cytoplasmic and nuclear localization **(Figure 3)**.

### The ORF2 protein is N-glycosylated on N1 and N3 sites but not on the N2 site

To characterize the impact of N-glycosylation site mutations on the ORF2 protein profile and oligomerization, ORF2 protein expression in supernatants and lysates of wt and mt PLC3/HEV-p6 cells was characterized by western blotting (WB). As shown in **Figure 4A**, a single band was detected in all cell lysates, which corresponds to ORF2i as previously described (19), immunofluorescence stainings observed in Figure 3 thus corresponded mainly to the subcellular localization of ORF2i proteins. We observed no shift of migration between wt and mutant proteins, even for the mutants carrying mutations that inhibit N-glycosylation. These results confirm that the ORF2i protein is not N-glycosylated, as previously described (19). In the supernatants of transfected cells, the three forms of the ORF2 protein (ORF2g, ORF2i and ORF2c) were identified (**Figure 4B**). Interestingly, we observed a shift of migration of ORF2g and ORF2c proteins for the mutants that inhibit N-glycosylation on N1 or N3 sites or both (N1S1, N1S2, N3S1, N3S2 and Gly(−)), as compared to wt protein (**Figure 4B** and **Figure 4D**). In contrast, no shift of migration was observed for proteins carrying mutations that do not affect glycosylation of N1 and N3 sites (N1S3, N3S3 and Gly(+)). These results indicate that both N1 and N3 sites of the ORF2 protein are likely occupied by N-glycans. Analyses of N2S1, N2S2 and N2S3 mutants showed that their ORF2g and ORF2c proteins behaved similarly whatever the introduced mutation (affecting or not glycosylation) prevented us from drawing conclusions about the N-glycosylation status of the N2 site. However, since it has been demonstrated that a proline residue right downstream of the N-X-S/T sequon constitutes an unfavorable context for N-linked glycan modification (21), we generated an additional mutant of the N2 site in which we replaced the Proline^313^ by an Alanine residue (N2S4) (**Figure 1**). Interestingly, N2S4 ORF2g and ORF2c proteins displayed a higher apparent molecular weight than the wt forms (**Figure 4B**), indicating that the N2S4 mutation likely leads to the addition of a supplementary N-glycan on ORF2 proteins. Together, these results indicate that the N2 site is likely not N-glycosylated in the context of wt ORF2 proteins.

**Figure 4:**
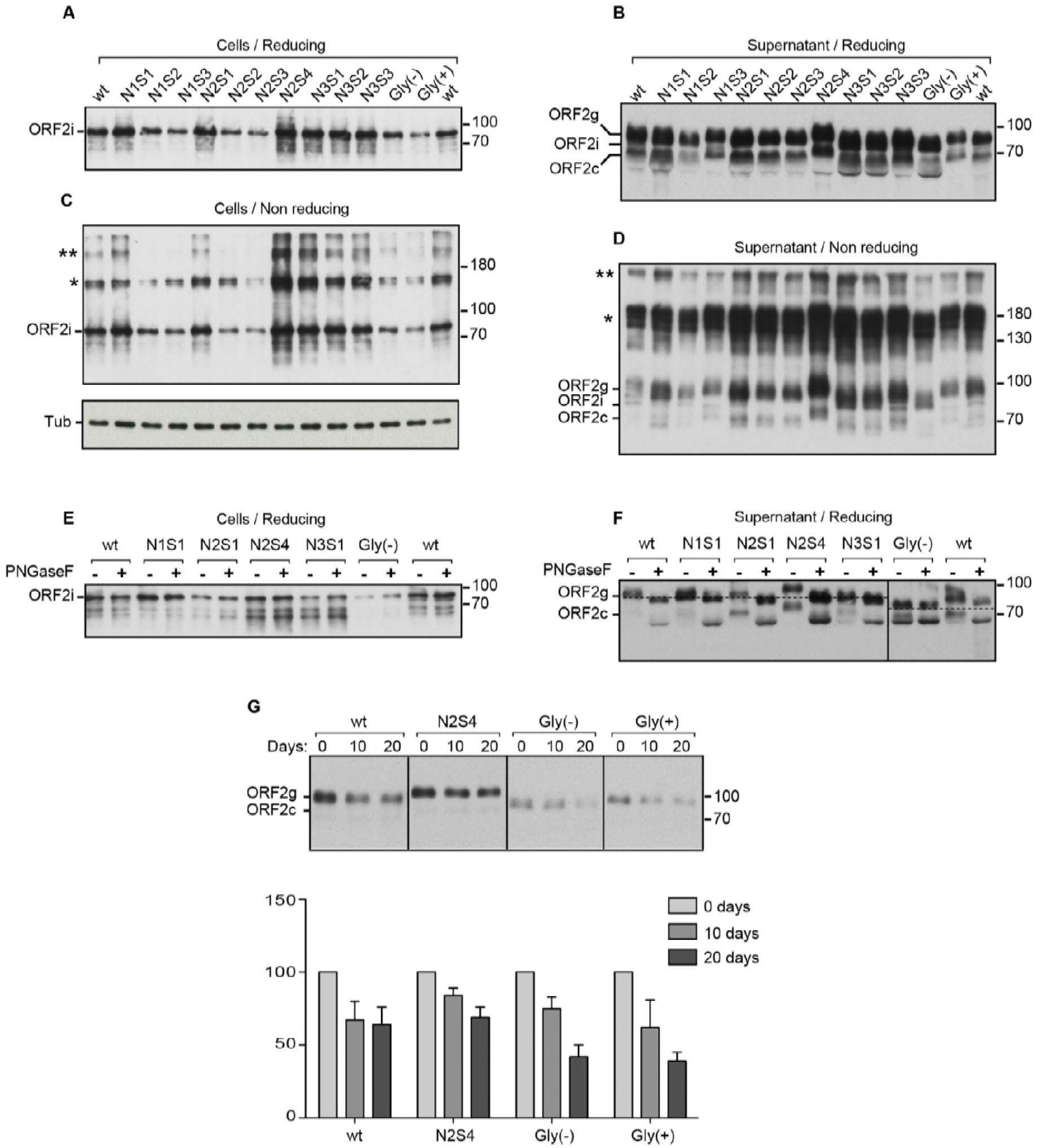
Characterization of wt and mutant ORF2 proteins by western blot (WB). Supernatants and lysates of wt and mt PLC3/HEV-p6 cells were collected at 10 days post-electroporation, normalized by protein quantification assay, and ORF2 protein was detected by WB using the 1E6 antibody. The representative results of two independent experiments are presented. Detection of ORF2 protein in cell lysates in reducing conditions (**A**) and in non-reducing conditions (**C**). Tubulin (Tub) was also detected to control protein loading. Detection of ORF2 protein in supernatants in reducing conditions (**B**) and in non-reducing conditions (**D**). Wt and mt PLC3/HEV-p6 cell lysates (**E**) and supernatants (**F**) were digested (+) or not (−) with Peptide-N-Glycosidase F (PNGaseF). (**G**) Supernatants of wt and mt PLC3/HEV-p6 cells were incubated for indicated times at 37°C. Relative ORF2 protein amounts were measured by densitometry. Values were adjusted to 100% for time 0 day. Results are from two independent experiments.

To further characterize which site of ORF2 protein is truly N-glycosylated, representative mutants were selected (N1S1, N2S1, N2S4, N3S1 and Gly(−)) and further characterized. Supernatants and lysates of wt and mt PLC3/HEV-p6 cells were denatured and digested with Peptide-N-Glycosidase F (PNGaseF), a glycosidase that cleaves between the innermost N-acetyl glucosamine and asparagine residues of N-glycoproteins. The proteins were resolved by SDS-PAGE and ORF2 proteins were detected by WB. As shown in **Figure 4E**, wt and mt ORF2i proteins expressed from cell lysates were resistant to glycosidase digestion, strengthening the fact that ORF2i protein in cell lysates is not N-glycosylated. In contrast, we observed a shift of migration between untreated (−) and treated (+) condition for the secreted wt and mutants N1S1, N2S1, N2S4 and N3S1 (**Figure 4F**). Importantly, no shift was observed for the Gly(−) double mutant carrying mutations that inhibit N-glycosylation on N1 and N3 sites, confirming that the N2 site is not occupied by N-glycans. In addition for the N2S4 mutant, we observed a higher migration shift between untreated (−) and treated (+) ORF2g and ORF2c proteins, as compared to wt (**Figure 4F**).

To further confirm which sites of N-glycosylation are occupied by glycans on ORF2 protein, we next performed mass spectrometry analyses. ORF2g/ORF2c proteins immunoprecipitated with an anti-ORF2 antibody (4B2) were treated or left untreated with PNGaseF. Proteins were resolved by SDS-PAGE and Colloïdal blue-stained bands corresponding to ORF2g and ORF2c proteins in WB (data not shown) were digested in-gel with trypsin (Tryp) or AspN and then analyzed by nano scale liquid chromatography coupled to tandem mass spectrometry. Detected peptides were compared to the theorical peptide sequences of non N-glycosylated or PNGaseF-deglycosylated ORF2 protein digested with Tryp or AspN. As shown in **Table 1**, when ORF2g/c proteins were not treated with PNGaseF, no peptides covering N1 and N3 sites were detected after digestion with Tryp or AspN. In contrast, upon treatment with PNGaseF, peptides covering N1 and N3 sites were detected after digestion with Tryp or AspN, indicating that N1 and N3 sites are most likely occupied by N-glycans (modifying the mass of peptides covering them and therefore their detection in the absence of PNGaseF treatment). Interestingly, peptides covering the N2 site were detected in both treated and untreated samples digested with Tryp or AspN, indicating that the probability that the site N2 is occupied by N-glycans is very low (**Table 1**).

**Table 1:** Mass spectrometry analyses of ORF2 N-glycosylation sites.

|  |  |  |  | Number of MSMS spectrum for ORF2g/ORF2c |  | Normalized total area of peptide peak for ORF2g/ORF2c |  | N-glycosylation probability |
| --- | --- | --- | --- | --- | --- | --- | --- | --- |
| Theoretical peptide sequences of non N-glycosylated or PNGaseF-deglycosylated ORF2 protein digested with Trypsin (Tryp) or AspN |  |  |  | Non treated | Treated by PNGaseF | Non treated | Treated by PNGaseF |  |
| Site N1 | Tryp | Non N-glycosylated<br>Deglycosylated by PNGaseF<sup>b</sup> | DVDSRGAIRRQY<u>N</u>LSTSP L TSSV ASGTNLVLYAAPLNPLLPLQDGTNTHIMATEASNYAQYR | ND<sup>a</sup> | ND | ND | ND | High |
|  |  |  | DVDSRGAIRRQY<u>D</u>LSTSP L TSSV ASGTNLVLYAAPLNPLLPLQDGTNTHIMATEASNYAQYR | ND | ND | ND | ND |  |
|  | AspN | Non N-glycosylated<br>Deglycosylated by PNGaseF<sup>c</sup> | DVDSRGAIRRQY<u>N</u>LSTSP L TSSV ASGTNLVLYAAPLNPLLPLQ | ND | ND | ND | ND |  |
|  |  |  | DVDSRGAIRRQY | ND | 2/2 | 0.0/0.0% | 0.5/1.2% |  |
| Site N2 | Tryp | Non N-glycosylated<br>Deglycosylated by PNGaseF | DFALELEFR<u>N</u>LTPGNTNT | 9/6 | 9/8 | 6.8/4.6% | 5.7/4.1% | Low |
|  |  |  | DFALELEFR<u>D</u>LTPGNTNT | ND | ND | ND | ND |  |
|  | AspN | Non N-glycosylated<br>Deglycosylated by PNGaseF | DFALELEFR<u>N</u>LTPGNTNTRVSRYTSTAQHRLRRGA | ND | ND | ND | ND |  |
|  |  |  | DFALELEFR | 2/2 | 6/6 | 0.1/0.2% | 2.4/2.3% |  |
| Site N3 | Tryp | Non N-glycosylated<br>Deglycosylated by PNGaseF | EAGTTRAGYPYNY<u>N</u>TTASDQILIENAAGHR | ND | ND | ND | ND | High |
|  |  |  | EAGTTRAGYPYNY<u>D</u>TTASDQILIENAAGHR | ND | 13/8 | 0.1/0.0% | 5.6/7.4% |  |
|  | AspN | Non N-glycosylated<br>Deglycosylated by PNGaseF | EAGTTRAGYPYNY<u>N</u>TTAS | ND | ND |  |  |  |
|  |  |  | EAGTTRAGYPYNY | ND | 2/2 | 0.0/0.0% | 1.1/0.8% |  |

<sup>a</sup> Not detected
<sup>b</sup> Upon PNGaseF treatment, the asparagine (N) residues from which the glycans have been removed are deaminated to aspartic acid (D) residues.
<sup>c</sup> Deamination of glycosylated N residues to D residues by PNGaseF treatment generates cleavage sites for AspN

Taken together, our results demonstrate that the ORF2 protein is N-glycosylated on N1 (^137^NLS) and N3 (^562^NTT) sites but not on the N2 (^310^NLT) site.

### Impact of mutations of ORF2 N-glycosylation sites on the oligomerization and stability of the ORF2 capsid protein

The impact of mutations of ORF2 N-glycosylation sites on the oligomerization of the ORF2 capsid protein was next assessed by analyzing supernatants and lysates of wt and mt PLC3/HEV-p6 cells prepared in non-reducing conditions. Oligomers of ORF2i and glycosylated ORF2 proteins were clearly detected (**Figure 4C** and **Figure 4D**, asterisks). Taking into account the differences in individual protein expression levels (**Figure 4A**), no significant differences in the oligomerization levels of ORF2 proteins was observed between wt and mutants.

We next analyzed the significance of ORF2 N-glycosylation for protein stability. For this purpose, supernatants of wt and mt PLC3/HEV-p6 cells were kept during 0, 10 and 20 days at 37°C and analyzed by WB (**Figure 4G**). Surprisingly, even after 20 days of incubation, ORF2g and ORF2c proteins were still readily detected, indicating that these proteins are very stable and poorly degraded in culture medium. Quantification by densitometry of ORF2 levels in wt and N2S4 supernatants, showed that only 30-35% of proteins were degraded after 20 days. In contrast, 60% of Gly(−) and Gly(+) proteins were degraded after 20 days, indicating that these mutants are less stable in culture medium but in a glycosylation-independent manner.

### Mutations of N-glycosylation sites do not modify ORF2 antibody recognition

To determine the impact of mutations of N-glycosylation sites on the anti-ORF2 antibody recognition, ORF2 proteins in cell lysates and supernatants of wt and mt PLC3/HEV-p6 cells were immunoprecipitated with either a linear (1E6) or two conformational (4B2 and 2E2, (22)) anti-ORF2 MAbs, and analyzed by WB (**Figure 5**). Although some detection differences were observed for several mutants, likely reflecting some differences in individual protein expression levels (**Figure 5A**, input), mutant ORF2i proteins were recognized by the three antibodies (**Figure 5A**). The supernatants were standardized according to the ORF2i expression levels and then immunoprecipitated with 1E6, 4B2 and 2E2 MAbs. As shown in **Figure 5B**, ORF2g and ORF2c proteins were equally immunoprecipitated by the three antibodies. We also quantified levels of secreted ORF2 proteins with the Wantaï HEV-antigen ELISA^Plus^ assay that has been recently marketed for HEV diagnosis and that works with monoclonal and polyclonal antibodies (Wantaï Biological Pharmacy Enterprise). As shown in **Figure 5C**, no significant differences were observed in protein detection between wt and mt ORF2 proteins.

**Figure 5:**
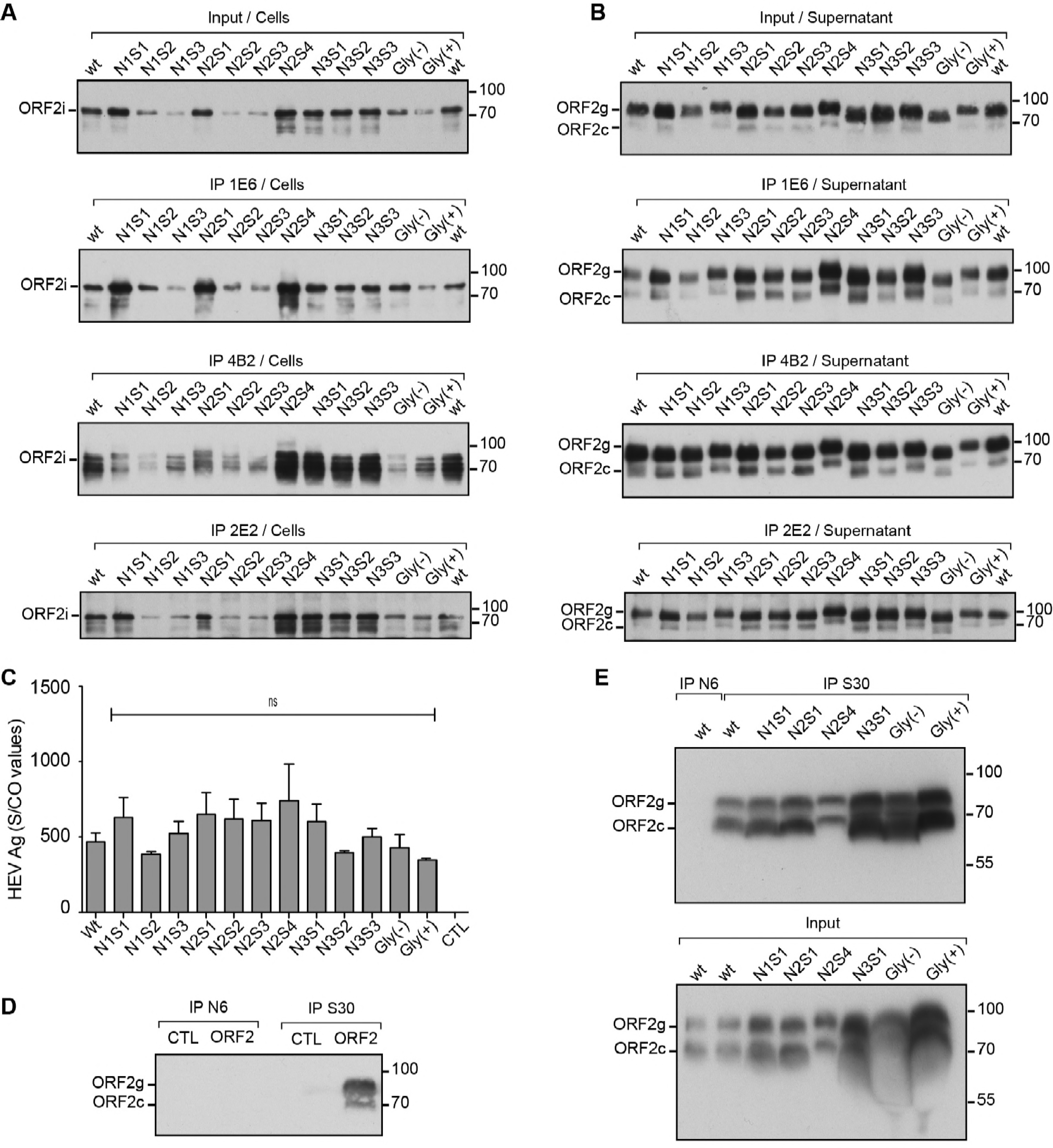
Impact of mutations of ORF2 protein N-glycosylation sites on antibody recognition. (**A** and **B**) Supernatants and lysates of wt and mt PLC3/HEV-p6 cells were collected at 10 days post-electroporation. Proteins in cell lysates and supernatants were normalized by protein quantification assay on cell lysates. ORF2 proteins were immunoprecipated (IP) by using the 1E6 linear anti-ORF2 antibody or the 4B2 and 2E2 conformational anti-ORF2 antibodies, as indicated. Input of ORF2 proteins used for immunoprecipitations are shown. ORF2 proteins were detected by WB using 1E6 antibody. **(C)** Detection of HEV-Ag in supernatants using the Wantaï HEV-Ag ELISA^Plus^ kit. Results are presented as signal to cut-off ratios (S/CO). (**D**) A serum from a non-infected patient (N6) and a serum from a patient who has cleared his HEV infection (S30) were both incubated with protein A-agarose beads and then with ORF2g/ORF2c proteins (ORF2) or PBS (CTL). ORF2 proteins were next detected by WB using 1E6 antibody. ORF2g/ORF2c proteins were isolated on iodixanol cushions (**E**) (Top) ORF2g/c proteins isolated from supernatants of wt and mt PLC3/HEV-p6 cells were immunoprecipitated with S30-immobilized beads. ORF2g/c proteins from supernatant of wt PLC3/HEV-p6 cells immunoprecipitated with N6-immobilized beads were used as a control. (Bottom) Input of ORF2g/ORF2c proteins used for immunoprecipitations are shown. Representative results of two independent experiments are shown.

We next sought to determine whether mutations of N-glycosylation sites affect recognition by patient antibodies. For this purpose, we used a serum from a patient who has cleared his HEV infection (serum S30) and a serum from a non-infected patient (serum N6). Sera were incubated with protein A-agarose beads and then with ORF2g/ORF2c proteins or PBS (CTL) (**Figure 5D**). ORF2g/ORF2c proteins were isolated on iodixanol cushions, as previously described (19). ORF2g and ORF2c proteins were specifically immunoprecipitated by immunoglobulins raised in the S30 patient serum whereas no proteins were precipitated by the N6 negative serum. As shown in **Figure 5E**, wt and mt ORF2g/ORF2c proteins were all recognized by antibodies from the S30 patient serum.

Taken together, these results demonstrate that mutations of ORF2 N-glycosylation sites do not modify the recognition of ORF2 protein by linear, conformational and patient antibodies. Thus, these mutations induce no major changes in ORF2 protein folding and N-glycosylation of ORF2 protein does not play a major role in antibody recognition. Importantly, our results show that ORF2g and ORF2c proteins are highly recognized by patient antibodies.

### The ORF2 protein translocates into the nucleus of infected cells independently of its N-glycosylation status

Since a recent study (20) and our confocal microscopy analyses (**Figure 3**) suggest that, in addition to its cytoplasmic localization, the ORF2 protein might also translocate into the nucleus of infected cells, we next further characterized this process by WB and immunofluorescence. We prepared cytoplasmic and nuclear extracts from wt and mt PLC3/HEV-p6 cells. Proteins were resolved by SDS-PAGE and the ORF2 protein was detected by WB. Cytoplasmic-specific anti-tubulin and nuclear envelope-specific anti-Lamin-B1 antibodies were used to control the quality of extractions. As expected, the ORF2i protein was detected in cytoplasmic extracts of wt and mt PLC3/HEV-p6 cells (**Figure 6A**). Interestingly, wt and mutant ORF2 proteins were also detected in nuclear extracts (**Figure 6B**). We named this protein ORF2ni for nuclear ORF2i. Among mutants of N-glycosylation, some differences in the ORF2ni detection were observed (**Figure 6B**). N1S3, N2S2, N2S3, Gly(−) and Gly(+) mutants showed reduced ORF2ni amounts whereas nuclear translocation was likely not affected for the other mutants, as compared to the wt protein. In order to quantify the effect of mutations of N-glycosylation sites on nuclear translocation, nuclear fluorescence intensity of ORF2 protein was measured on 50 cells for each mutant with the ImageJ software. As shown in **Figure 6C**, a significant reduction of nuclear translocation was observed for all mutants excepted for the N2S1 mutant and the three mutants of the N3 site. However, we did not observe any correlation between the status of N-glycosylation of ORF2 and its nuclear localization.

**Figure 6:**
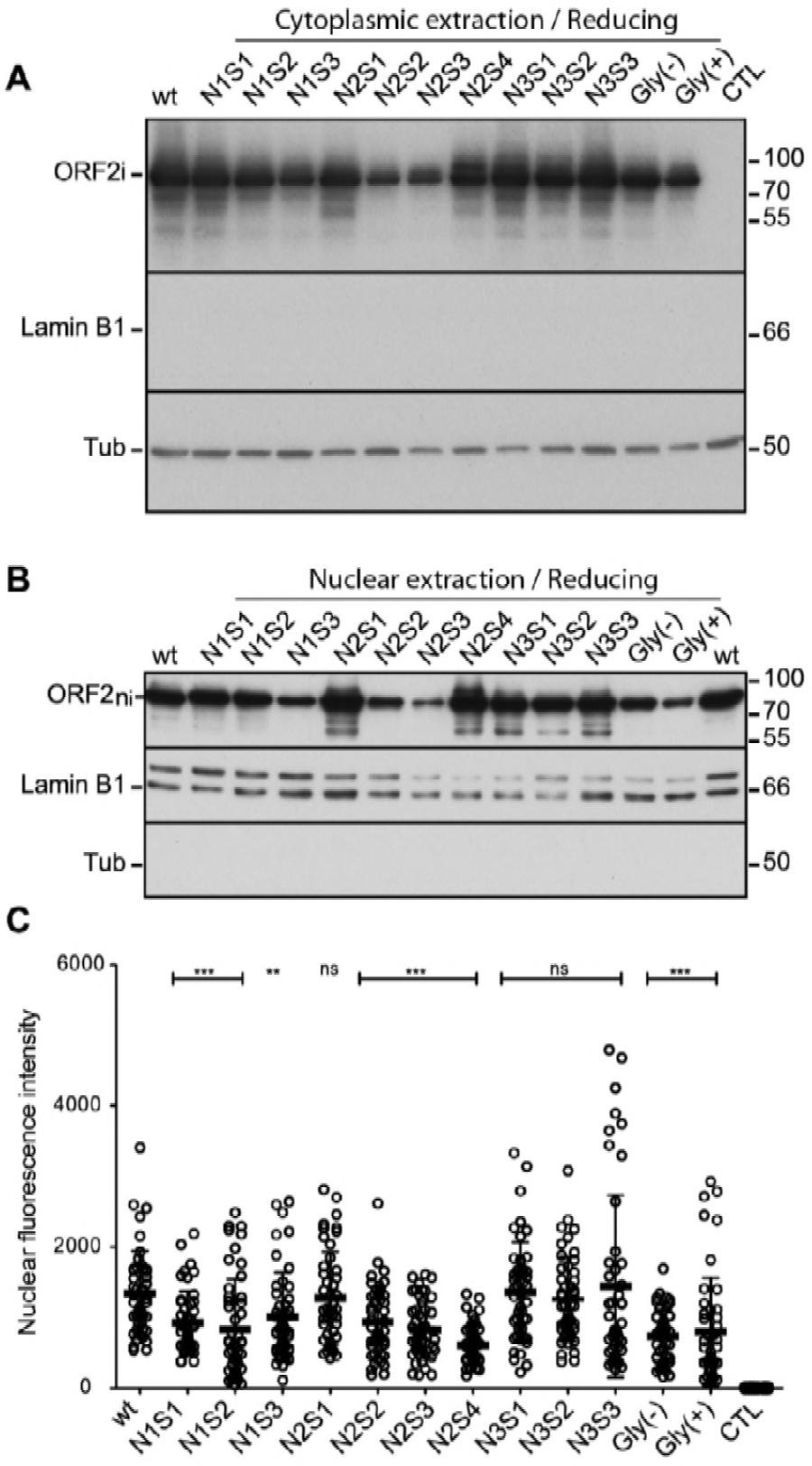
Impact of mutations of N-glycosylation sites on ORF2 protein nuclear localization. Cytoplasmic (**A**) and nuclear (**B**) extracts from wt and mt PLC3/HEV-p6 cells were prepared with the NE-PER Nuclear and Cytoplasmic extraction Kit. ORF2i and ORF2ni (nuclear ORF2i) proteins were detected by WB. LaminB1 and tubulin (Tub) were also detected to evaluate the quality of extractions. Representative results of two independent experiments are shown. (**C**) wt and mt PLC3/HEV-p6 cells were fixed at 3 d.p.e., and ORF2 protein was stained by using the 1E6 antibody. Cells (n=50) were analysed by LSM 800 confocal laser-scanning (Zeiss) using x40 oil immersion lens. The nuclear fluorescence intensity of ORF2 protein was determined using the ImageJ software.

Altogether, our results demonstrate that, during HEV lifecycle, the ORF2 capsid protein is translocated into the nucleus of infected cells. This nuclear localization is not closely related to the ORF2 protein N-glycosylation.

### Impact of mutations of ORF2 protein N-glycosylation sites on viral assembly and infectivity

Next, we analyzed the impact of mutations of ORF2 protein N-glycosylation sites on viral RNA production and infectivity. Supernatants of wt and mt PLC3/HEV-p6 cells were collected at 2, 6 and 10 d.p.e. and then processed for RNA level quantification by RT-qPCR (**Figure 7A**) and determination of extracellular infectious titers (**Figure 7B**). It has to be noted that, in order to specifically quantify capsid-protected RNA genomes, extractions and quantifications of extracellular RNA were performed after treatment of supernatants with RNase. Quantification of extracellular RNA genomes and calculation of fold increase between 2, 6 and 10 d.p.e., which are in line with HEV RNA replication and secretion of capsid-protected RNA genomes, showed that mutants of the N2 site led to a reduction of RNA replication and/or secretion of capsid-protected RNA genomes (**Figure 7A**), whereas extracellular RNA levels of N1, N3 and double mutants were similar to wt genome. To determine viral infectivity, Huh-7.5 cells were infected with serial dilutions of supernatants and processed for ORF2 staining at 3 d.p.i. Viral titers were determined by quantifying focus forming units (FFU/mL) (**Figure 7B**). As expected, supernatants of N2 mutants were not infectious. Supernatants of N1 and double mutants displayed reduced infectivity. Although the N3S1 mutant was slightly less infectious, other mutants of the N3 site displayed infectious titers similar to wt strain. These results indicate that mutations of the N1 site inhibit infectivity of viral particles whereas mutations of the N3 site have no major impact on the biogenesis of infectious HEV particles.

**Figure 7:**
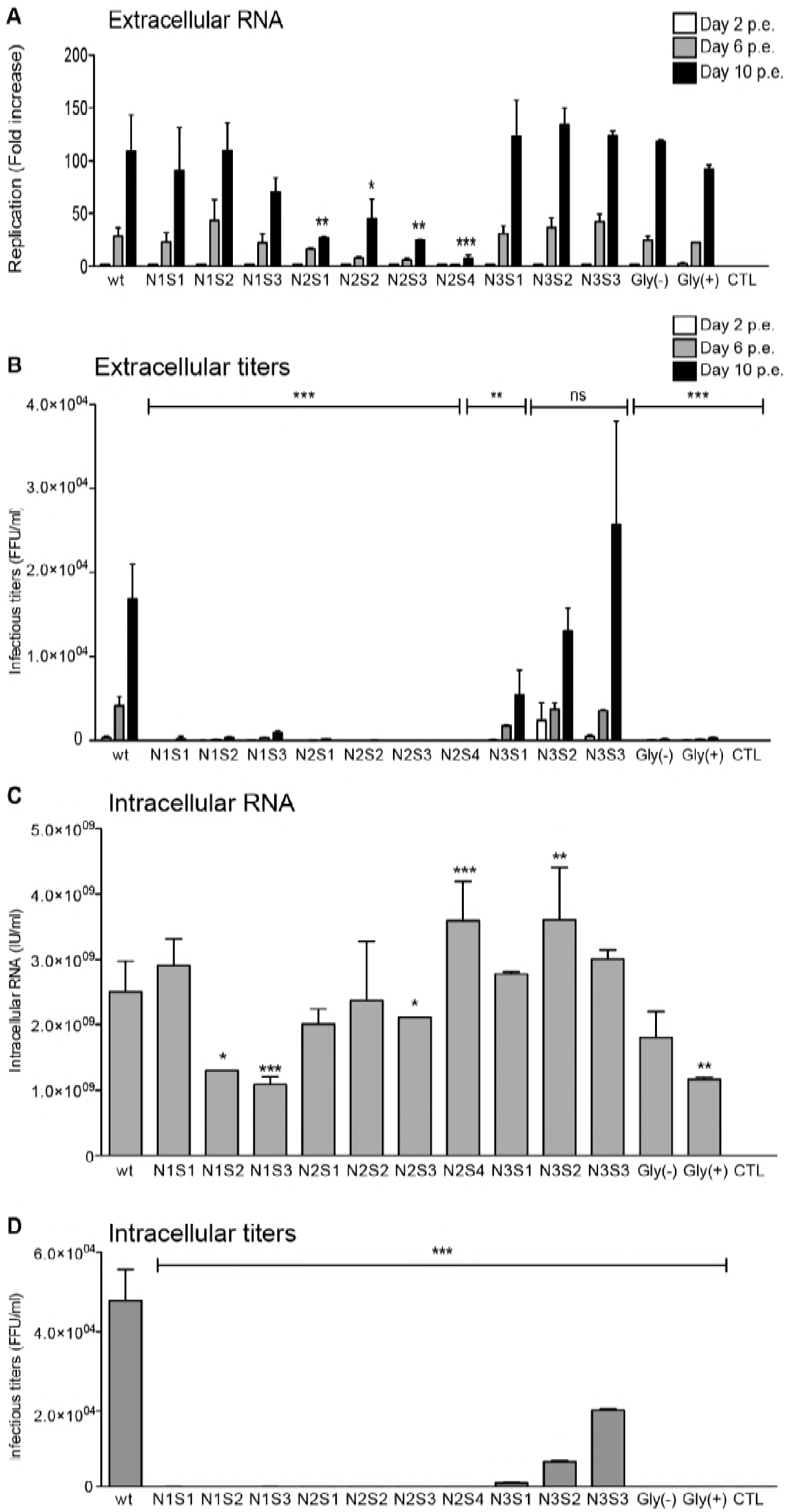
Impact of mutations of ORF2 protein N-glycosylation sites on viral assembly and infectivity. **(A)** The level of HEV RNAs in the supernatants of wt and mt PLC3/HEV-p6 cells collected at 2, 6 and 10 d.p.e. was measured by RT-qPCR (IU/ml). For each condition, values are presented as fold increase compared to RNA level measured at 2 d.p.e. Presented data are the mean of two independent experiments performed in duplicate **(B)** Naïve Huh.7.5 cells were infected with serial dilutions of supernatants collected at 2, 6 and 10 d.p.e. At 3 days post-infection, cells were fixed and ORF2 protein was detected by immunofluorescence. Focus forming Units (FFU) were determined and the results are presented as FFU/mL. Presented data are the mean of two independent experiments performed in triplicate. **(C)** Intracellular viral particles were extracted at 10 d.p.e. and the level of HEV RNAs was measured by RT-qPCR (IU/ml). Presented data are the mean of two independent experiments performed in duplicate. **(D)** An aliquot of intracellular viral particles was used to infect naïve Huh. 7.5 cells and the expression of ORF2 protein was analyzed by immunofluorescence. The Focus forming Unit (FFU) was determined and the results are presented as FFU/mL. The shown data are the mean of two independent experiments performed in triplicate.

In order to precisely define the impact of mutations on HEV RNA replication, we quantified the levels of intracellular viral RNA genomes. Although several significant differences were observed among mutants, intracellular RNA replication was globally not affected by the mutations of ORF2 protein N-glycosylation sites (**Figure 7C**). Finally, we determined the infectivity of intracellular particles produced in wt and mt PLC3/HEV-p6 cells (**Figure 7D**). N2S3, N2S4 and Gly(−) PLC3/HEV-p6 cells did not produce any infectious particles, indicating that these mutations are lethal for assembly of infectious HEV particles. N1 mutants, N2S1, N2S2 and Gly(+) mutants displayed highly reduced intracellular titers, indicating that these mutations affect assembly and infectivity of HEV particles. As for extracellular titers, the N3S1 mutant was slightly less infectious whereas N3S2 and N3S3 mutants displayed intracellular titers similar to wt strain.

Taken together, our results show that (i) mutations of N1 glycosylation site inhibit infectivity of HEV particles, (ii) mutations of N2 glycosylation site inhibit assembly of HEV particles and (iii) mutations of N3 glycosylation site have little or no effect on assembly of infectious HEV particles. Importantly, whatever the introduced mutation within the same site (affecting or not N-glycosylation) the same phenotype was observed indicating that N-glycosylation of ORF2 protein does not play any role in assembly of infectious HEV particles.

### Impact of mutations of ORF2 protein N-glycosylation sites on particle density

To further characterize our mutants, we produced large amounts of infectious supernatants by culturing wt and mt PLC3/HEV-p6 cells during 12 days. Supernatants were pooled, concentrated, and fractionated on an iodixanol gradient. ORF2 protein expression, density and RNA levels were determined for each fraction (**Figure 8 A-G**). As previously, only some representative mutants were analyzed (N1S1, N2S1, N2S4, N3S1, Gly(−) and Gly(+)). ORF2g and ORF2c proteins were highly enriched in fractions 4 and 5 whereas the ORF2i protein was mainly observed in fraction 5 or 6 of wt or mutants that assembled particles (N1S1, N3S1, Gly(−) and Gly(+)). In accordance with our previous results (**Figure 7**), N2S1 and N2S4 mutants displayed no ORF2i and no extracellular RNAs confirming that the mutations of N2 glycosylation site inhibit assembly of HEV particles. As described previously (19), only one major peak of RNA was detected in fraction 6, with a density of 1.11 g/mL, for wt and N3S1 gradients. In contrast, N1S1, Gly(−) and Gly(+) displayed a RNA peak in fraction 4 or 5 with a density of 1.10 g/mL, indicating that non-infectious mutants have a slightly lower density, as compared to infectious particles.

**Figure 8:**
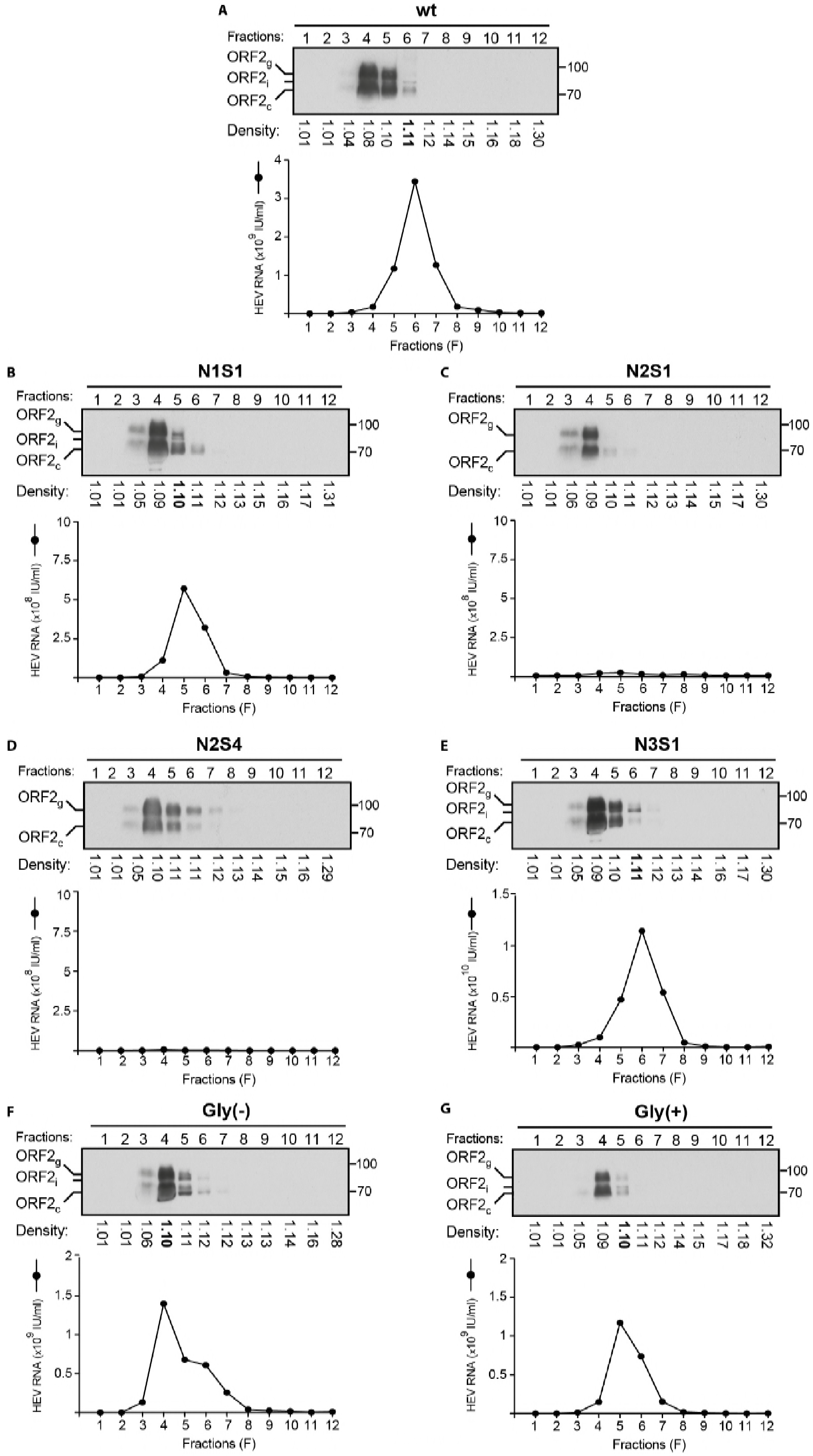
Impact of mutations of ORF2 protein N-glycosylation sites on particle density. Concentrated wt and mt PLC3/HEV-p6 cell supernatants were layered on an iodixanol gradient and ultracentrifuged. Twelve fractions were collected and their densities were measured. ORF2 expression was analyzed by WB using 1E6 antibody. HEV RNA levels in each fraction were quantified by RT-qPCR.

## Discussion

We recently demonstrated that during its infectious cycle, HEV produces three different forms of its ORF2 capsid protein (19). The ORF2i form (infectious / intracellular) is the form associated with the infectious particles. This protein of about 80 kDa is neither N-glycosylated or nor O-glycosylated. The ORF2g (glycosylated) and ORF2c (cleaved) forms, are proteins that are massively secreted but are not associated with infectious material. These two forms are approximately 90 kDa and 75 kDa, respectively. They are N-glycosylated, O-glycosylated and sialylated. In the present study, we identified the precise sequence of the ORF2c form that is likely a cleavage product of the ORF2g protein. Importantly, ORF2g/c proteins do not form particulate material but they are the major viral antigens present in the serum of HEV-infected patients. HEV may produce ORF2g/c proteins as immunological bait. Since these proteins are likely extremely stable in infected patients (23), they might represent markers of the evolution of hepatitis E infection. It is therefore essential to determine the molecular and cellular mechanisms leading to the biogenesis of the different forms of ORF2 protein. In the present study, we analyzed the N-glycosylation status of ORF2 proteins and the biological relevance of this post-translational modification in the HEV life cycle. We also obtained new insights into the molecular mechanisms of ORF2 expression.

The ORF2 protein has been largely studied and is the most characterized HEV viral protein. Notably, its glycosylation status has been analyzed by several laboratories but no clear evidence was depicted about the biological relevance of this post-translational modification in the HEV life cycle. As early as 1996, by using SV40-based expression vectors in COS-1 cells, Jameel and collaborators described the gt1 ORF2 protein as an 88kDa glycoprotein which is expressed intracellularly as well as on cell surface and has the potential to form non covalent homodimers (15). Later on, using the same expression system, they demonstrated that Asn-310 (N2 site) is the major but not the only site of N-glycan addition (14). In 2008, Graff et al. analyzed the impact of mutations within highly conserved glycosylation sites of the gt1 Sar55 infectious full-lenght genome (13). They observed no difference in electrophoretic migration rates of wildtype glycosylated and mutant nonglycosylated ORF2 proteins in cell lysates, suggesting that ORF2 protein accumulating within transfected S10-3 cells is not glycosylated. They also demonstrated that mutation of the first 2 glycosylation sites prevented virion assembly and mutation of the third site did not affect particle formation and RNA encapsidation but the particles were not infectious. However, conservative mutations that did not affect glycosylation also prevented infection, indicating that mutations of N-glycosylation sites were lethal because they perturbed protein structure rather than because they eliminated glycosylation (13). In our study, by using site-directed mutagenesis of the three N-glycosylation sites, PNGaseF treatment and mass spectrometry analyses of ORF2 protein expressed from our efficient gt3 p6 full-length strain-based HEV cell culture system, we robustly demonstrated that the ORF2 protein is N-glycosylated on N1 (^137^NLS) and N3 (^562^NTT) sites but not on the N2 (^310^NLT) site. The absence of N-glycan addition on N2 site is likely due to the presence of a Proline^313^ residue immediately downstream of the ^310^NLT sequon, which has a deleterious effect on N-glycan addition (24). In accordance with this claim, we showed that the Proline^313^ to Alanine^313^ substitution (N2S4 mutant) induced additional ORF2 N-glycosylation. It has to be noted that alignment of 293 sequences from genotype 1 to genotype 4 HEV strains showed that the Proline^313^ is highly conserved (25), suggesting that the absence of N-glycans on ORF2 N2 site is likely a conserved process among HEV genotypes. We have been able to robustly analyze the glycosylation status of ORF2 thanks to the high expression levels of this protein in our replicative system. It would be now also interesting to develop a similar cell culture system for the gt1 HEV in order to robustly analyzed the glycosylation status of the gt1 ORF2 protein.

We performed an extensive characterization of ORF2 mutants in which the three potential N-glycosylation sites were mutated individually or in combination. Whatever the introduced mutations, we observed no significant differences in expression, secretion, oligomerization, stability and recognition by antibodies of the secreted ORF2g and ORF2c proteins. However, we showed that ORF2g and ORF2c proteins were very stable and poorly degraded when kept in culture medium at 37°C, which is consistent with previous observations showing that ORF2 protein remained detectable for more than 100 days after HEV RNA clearance in ribavirin-treated patients with chronic HEV infection (23). Importantly, we showed that ORF2g and ORF2c proteins were the main antigens recognized by antibodies from a patient who had cleared his HEV infection (serum S30), indicating that these glycosylated ORF2 forms are the main targets of the humoral response during HEV infection.

The level of ORF2i protein expression and its recognition by MAbs were not affected by mutations of N-glycosylation sites, which is in line with the fact that this protein is not glycosylated. However, during our study, we demonstrated that intracellular ORF2 proteins displayed both cytoplasmic and nuclear localization, as recently observed by immunohistochemistry of liver biopsy from patients with hepatitis E (20). By performing differential extractions and detection by western blotting, we demonstrated that the ORF2i protein localized in the nuclear fraction of HEV producing cells. We named this fraction of ORF2i, ORF2ni for nuclear ORF2i protein. Characterization of N-glycosylation mutants showed that nuclear localization of ORF2ni is not closely related to the ORF2 protein N-glycosylation. Altogether, our results demonstrate that, during HEV life cycle, the ORF2 capsid protein is translocated into the nucleus of infected cells to presumably control certain cellular functions to promote viral replication and/or alter the antiviral response of the infected cell. Further studies are now necessary to decipher the mechanisms and significance of ORF2 nuclear translocation in the HEV life cycle.

We analyzed the impact of mutations of ORF2 protein N-glycosylation sites on viral RNA production and particle infectivity. None of the mutations had an effect on viral replication. On the other hand, mutations of N1 glycosylation site inhibited infectivity of HEV particles, mutations of N2 glycosylation site abolished assembly of HEV particles and mutations of N3 glycosylation site had little or no effect on assembly of infectious HEV particles. Importantly, as observed by Graff et al. (13), whatever the introduced mutation, affecting or not N-glycosylation, the same effects were observed, indicating that N-glycosylation of ORF2 protein does not play any role in assembly of infectious HEV particles. Mutations likely induce modifications in ORF2i protein domains involved in particle assembly. Interestingly, we observed that non-infectious particles were characterized by a slightly lower density, as compared to infectious particles, which might reflect subtle differences in ORF2 assembly leading to non-infectious particles. It has been suggested that cytoplasmic localization of the ORF2 protein would depend on its ability to be retrotranslocated from the ER by a glycosylation-dependent process (26). In that case, a non-glycosylated form of ORF2, such as our Gly(−) mutant, will not generate any cytoplasmic ORF2i protein. However, the Gly(−) mutant produced ORF2i protein levels similar to that of the wt ORF2, indicating that ORF2i proteins are likely not generated from a glycosylation-dependent retrotranslocation process.

Since the ORF2g/c proteins might interfere with the capacity of HEV virions to infect target cells (19), we also performed inhibition assays of HEV infection in the presence of mutant ORF2g/c proteins. However, we could not make any correlation between glycosylation status and inhibition levels (data not shown).

Altogether, our results demonstrate that ORF2 N-glycosylation is not essential in the HEV life cycle. However, the ORF2g/c proteins are also modified by O-glycosylation and sialic acids, as recently demonstrated (19, 27). Further studies are now necessary to identify the significance of these modifications in the functionality of ORF2g/c proteins in HEV infection. Anyway, our findings strengthen our hypothesis that the HEV life cycle or at least the production of viral particles might be tightly regulated by the differential addressing of the ORF2 capsid protein. Indeed, the identification of the 3 forms of ORF2 led us to suggest the existence of two production pathways for the HEV capsid protein: (i) a major non-productive pathway in which ORF2 proteins are addressed to the secretion route where they are glycosylated, maturated and quickly secreted. (ii) a productive pathway in which cytosolic ORF2 proteins are delivered to the virion assembly sites (19). The identification of the nuclear ORF2ni protein suggests now that a fine balance of ORF2 addressing likely occurs between cytosolic, nuclear and reticular pathways.

## Materials and Methods

### Chemicals and cell cultures

PLC3 (19) and Huh-7.5 (28) cells were grown in Dulbecco’s modified Eagle’s medium (DMEM) supplemented with 10% inactivated fetal calf serum and 1% of Non-Essential amino acids (Life Technologies) at 37°C. Transfected PLC3 cells were maintained at 32°C in a medium containing DMEM/M199 (1v:1v), 1mg/ml of lipid-rich albumin (Albumax I™), 1% of Non-Essential amino acids and 1% of pyruvate sodium (Life Technologies).

### Plasmids and transfection

The plasmid pBlueScript SK(+) carrying the DNA of the full length genome of adapted gt3 Kernow C-1 strain, (HEV-p6, GenBank accession number JQ679013, kindly provided by S.U Emerson) was used as a template (18). The mutants of the ORF2 N-glycosylation sites were generated by site directed mutagenesis. Individual mutations were introduced by sequential PCR steps, as described previously (29), using the Q5 High-Fidelity 2X Master Mix (New England Biolabs, NEB), then digestions with restriction enzymes and ligation were performed. The double mutants of N-glycosylation sites were generated by exchanging the mutant fragments from single mutants using specific restriction sites. All the mutations were verified by DNA sequencing.

To prepare genomic HEV RNAs (capped RNA), the wild type (wt) and mutant (mt) pBlueScript SK(+) HEV-p6 plasmids were linearized at their 3’ end with the MluI restriction enzyme (NEB) and transcribed with the mMESSAGE mMACHINE^®^ kit (Ambion). Capped RNAs were next delivered to PLC3 cells by electroporation using a Gene Pulser Xcell™ apparatus (Bio-Rad) (19).

### Patient samples

Patient samples were collected in France between 2014 and 2016. Samples were obtained only *via* standard viral diagnostics following a physician’s order (no supplemental or modified sampling). Data were analyzed anonymously. The French Public Health Law (CSP Art L 1121-1.1) does not require written informed consent from patients for such a protocol.

### Kinetic experiments and virus production

PLC3 cells were electroporated with wild type and mutant HEV-p6 RNAs (20μg/3×10^6^ cells). For kinetics experiments, supernatants were harvested 2, 6 and 10 days post-electroporation (d.p.e) and then used for viral titers, RNAs quantification and WB analysis. Transfected cells (6×10^6^) were lysed 10 d.p.e. in buffer containing 10mM TrisHCl (pH 7), 150mM NaCl, 2mM EDTA, 0.5% Triton X-100, 1mM PMSF and protease inhibitor cocktail (Complete; Roche). Supernatants and cell lysates were stored at −80°C until analysis.

### Antibodies

Three mouse anti-HEV ORF2 monoclonal antibodies (MAb) were used: (i) the linear 1E6 anti-ORF2 MAb (antibody registry #AB-827236, Millipore), (ii) the conformational 4B2 anti-ORF2 MAb (antibody registry #AB-571018, Millipore) and (iii) the conformational 2E2 anti-ORF2 MAb (antibody registry #AB-571017, Millipore). Mouse anti-β tubulin (antibody registry #AB-609915) was from Sigma and rabbit anti-Lamin B1 (antibody registry #AB-443298) antibody was from Abcam. Secondary antibodies were purchased from Jackson ImmunoResearch.

### Indirect immunofluorescence

PLC3 cells electroporated with wildtype and mutants of HEV-p6 RNAs (wt and mt PLC3/HEV-p6 cells) were grown on coverslips in 24-well plates and fixed 3 d.p.e. with 3% of Paraformaldehyde (PFA). After 20 minutes (min), cells were washed twice with phosphate-buffered saline (PBS) and permeabilized for 5 min with cold methanol and then with 0.5% Triton X-100 for 30 min. Cells were incubated in PBS containing 10% goat serum for 30 min at room temperature (RT) and stained with the 1E6 anti-ORF2 MAb for 30 min at RT followed by a Cy3-conjugated goat anti-mouse antibody for 20 min at RT. The nuclei were stained with DAPI (4’,6-diamidino-2-phenylindole). After 2 washes with PBS, coverslips were mounted with Mowiol 4-88 (Calbiochem) on glass slides and analyzed with a LSM 800 confocal laser-scanning microscope (Zeiss) using a x40/1.4 numerical aperture oil immersion lens.

### Quantification of the ORF2 protein nuclear fluorescence

The method was adapted from McCloy et al. (30). Briefly, the ORF2 protein nuclear fluorescence was determined using ImageJ software (version 1.51, NIH). The regions of interest (ROI) were drawn around the nuclei on the immunofluorescence image from PLC3/HEV-p6 wt and mt using imageJ ROI tools. Area, integrated density and mean gray values were measured. Then, corrected total cell fluorescence (CTCF) was calculated by the following formula: CTCF= integrated density – (area of selected electroporated cells x mean of background fluorescence around the cells). The exact nuclear fluorescence was = CTCF-the mean of the integrated density of non-infected cells.

### Western blotting analyses

Western blotting analyses were performed as described previously (31). Briefly, supernatants and lysates of wt and mt PLC3/HEV-p6 cells were heated for 20 min at 80°C in the presence of Laemmli buffer (reducing or non-reducing). Samples were then separated by SDS-PAGE and transferred onto nitrocellulose membranes (Hybond-ECL, Amersham). The targeted proteins were detected with specific antibodies and corresponding peroxidase-conjugated secondary antibodies. The detection of proteins was done by chemiluminescence analysis (ECL, Amersham).

### PNGase-F treatment

Supernatants and lysates of wt and mt PLC3/HEV-p6 cells were denaturated for 10min at 95°C in glycoprotein denaturing buffer (NEB). Digestions with Peptide-N-Glycosidase F (PNGaseF, NEB) were carried out for 4h at 37°C in the presence of 1% NP40 and the buffer provided by the manufacturer (NEB). Samples prepared in the same conditions but without glycosidase were used as controls.

### Nuclear and cytoplasmic extractions

Confluent T75 flasks of wt and mt PLC3/HEV-p6 cells (6 × 10^6^ cells) were harvested 12 d.p.e. with trypsin-EDTA. Cells were centrifuged at 4000rpm for 5 min and washed thrice with PBS. Nuclear and cytoplasmic proteins were extracted using the NE-PER Nuclear and Cytoplasmic extraction Kit (Thermo scientific) following the manufacturer’s recommendations.

### Immunoprecipitations

Polyclonal rabbit anti-mouse antibody (DAKO) was bound to protein A-agarose beads and incubated overnight with mouse anti-ORF2 MAb (1E6 or 4B2). Beads were washed thrice with PBS and then incubated for 2 hours at room temperature with supernatants or lysates of wt and mt PLC3/HEV-p6 cells. Beads were washed six times with PBS 0.5% NP40 and then heated at 80°C for 20 min in Laemmli buffer. Proteins were separated by SDS-PAGE and ORF2 proteins were detected by WB using the 1E6 MAb.

### Quantification of the ORF2 protein levels by ELISA

The supernatant of wt and mt PLC3/HEV-p6 cells and non-transfected PLC3 cells (CTL) were diluted in PBS (1:250 and 1:500). HEV ORF2 Ag levels were measured with the Wantaï HEV-Ag ELISA^Plus^ kit (Wantaï Biological Pharmacy Enterprise), as recommended by the manufacturer.

### Intracellular viral particles and RNAs

The procedure was adapted from Emerson et al. (32). Briefly, confluent T25 flasks of wt and mt PLC3/HEV-p6 cells were trypsinized and cells were centrifuged for 10 min at 1500 rpm. Cells were washed thrice with PBS. Intracellular viral particles were extracted by resuspending cells in 1ml of sterile MilliQ water at room temperature. Cells were vortexed vigorously for 20 min and then 110μl of sterile 10X PBS were added. Samples were clarified by centrifugation 2 min at 14000 rpm. The supernatants containing intracellular particles and RNAs were collected and stored at −80°C until analysis.

### HEV RNAs extraction and quantification

Supernatants collected at 2, 6 and 10 d.p.e and intracellular viral particles produced in wt and mt PLC3/HEV-p6 cells were submitted to viral RNAs extraction. HEV RNA levels were quantified by RT-qPCR, as described previously (33, 34).

### Infectious titers

Huh 7.5 cells (3 x 10^3^) seeded in 96-well plates the day before were infected with serial dilutions of supernatants or intracellular viral particles from wt and mt PLC3/HEV-p6 cells. Three days post-infection, cells were fixed and processed for indirect immunofluorescence. Cells labeled with anti-ORF2 antibody 1E6 were counted as infected cells. The number of infected cells was determined for each dilution and used to define the infectious titers in focus forming unit (FFU)/ml.

### Mass spectrometry

N-terminus identification of the ORF2c protein was performed as in Montpellier et al. (19). For N-glycans analyses, ORF2g/ORF2c proteins were immunoprecipitated with the 4B2 anti-ORF2 antibody and denaturated for 10min at 95°C in glycoprotein denaturing buffer (NEB). Proteins were treated or not with PNGaseF (19), as described above, and resolved by SDS-PAGE. Colloïdal blue stained bands corresponding to ORF2g and ORF2c proteins in WB were cut into two slices for in-gel digestion with trypsin or AspN. NanoLC-MSMS analyses of the protein digests were performed on a UltiMate-3000 RSLCnano System coupled to a Q-Exactive instrument (Thermo Fisher Scientific). Collected raw data were processed and converted into *.mgf peak list format with Proteome Discoverer 1.4 (Thermo Fisher Scientific). MS/MS data was interpreted using search engine Mascot (version 2.4.0, Matrix Science) with a tolerance on mass measurement of 10 ppm for precursor and 0.02 Da for fragment ions, against a composite targetdecoy database (40584 total entries) built with the sequences of ORF2 (H9E9C9_HEV) and the PNGaseF-deglycosylated ORF2 protein in which the three N-glycosylated sites were replaced by Asp(D) residues, fused with a Swissprot homo sapiens database (TaxID=9606, 20 May 2016, 20209 entries) and a list of classical contaminants (119 entries). Carbamidomethylation of cysteine residues, oxidation of methionine residues and protein N-terminal acetylation were searched as variable modifications. Up to three trypsin or Asp-N missed cleavage were allowed. Semi-specific cleavage was also authorized. Spectral counting was performed without MS score filtering. Peptides quantitation of ORF2 and PNGaseF-deglycosylated ORF2 protein was performed on MS1 level using Skyline (ver. 3.7) (35). After automated peak picking and retention time alignment of Skyline, a manual correction of wrong peak boundaries was performed and normalized total areas of peptide peaks were exported.

### Density gradients

Supernatants of wt and mt PLC3/HEV-p6 cells were collected and concentrated 100 times with Vivaspin ultrafiltration spin columns (Sartorius). Concentrated supernatants were layered on a 7.5-40% iodixanol gradient, which was centrifuged at 160,000g for 16h at 4°C. Twelve fractions of 1ml were collected and their density was measured by refractometry. Each fraction was used for RNA quantification and immunoblotting, as described above.

## Acknowledgments

This work was supported by the French “Agence Nationale de la Recherche sur le Sida et les hépatites virales” (ANRS). M. A. was supported by a fellowship from the ANRS. IMS was supported by PhD-grants from Ghent University and the Egyptian Government. PM was supported by an ‘Excellence of Science’ grant of the Research Foundation – Flanders (FWO-Vlaanderen) under grants n° EOS-30981113; and FWO project G0D2715N.

We thank Sophana Ung for his technical contribution. We thank Suzanne U. Emerson (NIH, USA) for providing us with reagents. We thank the imaging core facility of the BioImaging Center Lille-Nord de France for access to the instruments.

